# Impaired IFNγ responsiveness of monocyte-derived lung cells limits immunity to *Mycobacterium tuberculosis*

**DOI:** 10.1101/2025.10.06.680719

**Authors:** Weihao Zheng, Jason D. Limberis, Zachary P. Howard, Alexander Mohapatra, Enzo Takagi, Joel D. Ernst

**Affiliations:** Division of Experimental Medicine, Department of Medicine, University of California, San Francisco, California, USA; Division of Nephrology, Department of Medicine, University of California, San Francisco, California, USA

## Abstract

Lung mononuclear phagocyte (MNP) subsets differ in their ability to restrict *Mycobacterium tuberculosis* (Mtb) during chronic infection, yet the mechanisms underlying this difference are not well defined. Here, we show that CD11c^lo^ monocyte-derived cells (MNC1), the subset of lung cells that is most permissive for Mtb viability during chronic infection, express lower levels of interferon-gamma (IFNγ) signaling proteins, resulting in reduced responses to IFNγ compared to alveolar macrophages (AM) and CD11c^hi^ MNC2. Moreover, type I IFN signaling suppresses IFNγ-mediated MHC-II expression, impairing antigen-specific CD4 T cell activation by MNC1 cells. Importantly, prior immunity conferred by contained Mtb infection enhances IFNγ responsiveness of monocyte-derived cells, reducing bacterial burdens in lungs and within MNC subsets. Our findings indicate that heterogeneous IFNγ responsiveness is exploited by Mtb for persistence in vivo. Overcoming or bypassing impaired IFNγ responsiveness may guide the development of more effective TB vaccines and host-directed therapies.

## Introduction

Tuberculosis (TB), caused by the respiratory pathogen *Mycobacterium tuberculosis* (Mtb), returned to being the leading cause of death from a single infectious disease worldwide as the COVID-19 pandemic subsided. It is imperative to identify mechanisms of immune protection and pathogenesis in TB to facilitate the development of novel TB vaccines and therapies.

Mtb is known for its ability to evade elimination by immune responses and cause chronic infection^1–3^. Multiple studies, including recent single-cell RNA sequencing studies, have revealed significant heterogeneity among lung mononuclear phagocytes (MNP) during Mtb infection^4–6^. Moreover, mounting evidence has established that Mtb infects multiple lung MNP subsets, among them, monocyte-derived cells (MNC) serve as the major bacterial reservoir during chronic infection^4,7–10^. After development of adaptive immune responses, distinct lung MNP subsets differ in Mtb-killing ability. Specifically, alveolar macrophages (AM) have a superior ability to restrict Mtb compared to monocyte-derived MNC1 and MNC2 cells ^4,8^. Among these MNP subsets, CD11c^lo^ MNC1 is the most Mtb-permissive subset as defined by the number of live Mtb per infected cell, suggesting that functional heterogeneity of MNP contributes to Mtb persistence in vivo. We previously discovered heterogeneous lysosome function in these MNP subsets and found that in MNC1, lysosome deficiency is one cell type-specific host mechanism that contributes to Mtb persistence in vivo^8^. However, the mechanisms underlying the functional heterogeneity, particularly the differential ability of lung MNP subsets to respond to immune signals, remain incompletely defined.

Interferon-gamma (IFNγ) plays a critical role in host defense against Mtb, as evidenced by both human and murine studies^11–15^. Individuals with mutations in genes responsible for IFNγ immunity (e.g., *Ifng*, *Ifngr1*, *Ifngr2*, and *Stat1*) are susceptible to mycobacterial infections, including non-tuberculous mycobacteria and Mtb^11,16^. Studies utilizing knockout mice have demonstrated that both IFNγ and its receptor, IFNGR1, are essential for controlling Mtb infection, suggesting that the host’s ability to produce IFNγ or to respond to IFNγ is required for IFNγ-mediated protection against TB^12–14^. Both hematopoietic and nonhematopoietic IFNγ responsiveness are required for Mtb control^14^. Paradoxically, TB vaccines that predominantly induce IFNγ often fail to enhance lung protection^1^. Moreover, murine studies indicate that increasing IFNγ production does not enhance Mtb control in the lungs, suggesting that cellular responsiveness to IFNγ may be a key determinant of its efficacy ^17^. It remains unknown whether distinct lung MNP subsets differ in their ability to sense and transduce IFNγ signals during primary Mtb infection, and whether there are interventions that alter IFNγ responsiveness in lung MNP subsets to affect infection outcome. Elucidating cell-specific actions of IFNγ is crucial for a comprehensive understanding of IFNγ-mediated immunity in tuberculosis, to make it more effective.

In this study, we identify subset-specific differences in IFNγ responsiveness across distinct lung MNP subsets during Mtb infection. Specifically, compared with AM, Mtb-permissive MNC1 cells are less responsive to IFNγ and exhibit defective IFNγ signal transduction, while MNC2 cells exhibit intermediate responsiveness to IFNγ. Compared with AM and MNC2 cells, MNC1 cells express less MHC-II and have a reduced ability to present Mtb antigens to CD4 T cells, which is partially dependent on Type I IFN signaling. We further found that prior Mtb immunity can improve IFNγ responsiveness of MNC cells, leading to better Mtb control. Together, these findings provide mechanistic insights into the functional heterogeneity of lung MNP subsets during TB, suggesting that targeting IFNγ responsiveness may represent a strategy for developing novel TB therapies and vaccines.

## Results

### Differential responses to IFNγ in MNP subsets from *M. tuberculosis*-infected lungs

To investigate if lung MNP subsets might respond to IFNγ differently during chronic Mtb infection, we first analyzed our bulk RNA-seq data for three flow-sorted lung MNP subsets (AM, CD11c^lo^ MNC1, and CD11c^hi^ MNC2) from mice infected with Mtb for 28 days (GSE220147) (Figure S1)^8^. Gene Ontology (GO) analysis revealed that IFNγ-responsive genes were expressed at significantly lower levels in MNC1 compared to AM or MNC2 (Table S1). We further investigated the expression levels of IFNγ-responsive genes in a published gene list^6,18^. In this analysis, MNC1 cells exhibited decreased expression of IFNγ-responsive genes compared to AM and MNC2 (Figure 1A), suggesting that this lung cell subset responds to IFNγ less effectively. Since IFNγ receptors and downstream regulators/proteins in the pathway dictate the cellular response to IFNγ (Figure 1B), we investigated their expression levels. This revealed that MNC1 had lower mRNA expression of *Ifngr1, Ifngr2*, *Stat1*, *Irf1*, *Cxcl9*, and *Nos2* compared to MNC2, while AM expressed significantly more *Ifngr2* mRNA than MNC1, with a similar but non-significant trend observed for *Ifngr1, Stat1*, and *Cxcl9* (Figure 1C). Using flow cytometry, we confirmed that MNC1 expressed lower levels of IFNGR2, STAT1, and IRF1 proteins, as well as the IFNγ-responsive proteins CXCL9 and NOS2, compared to AM or MNC2 (Figure 1D and Figure S2). Although AM expressed more IFNGR1 protein than MNC1, the protein level of IFNGR1 did not differ between MNC1 and MNC2 (Figure 1D and Figure S2). Together, these findings indicate that there are differential signaling and responses to IFNγ in these lung MNP subsets.

**Figure 1.**
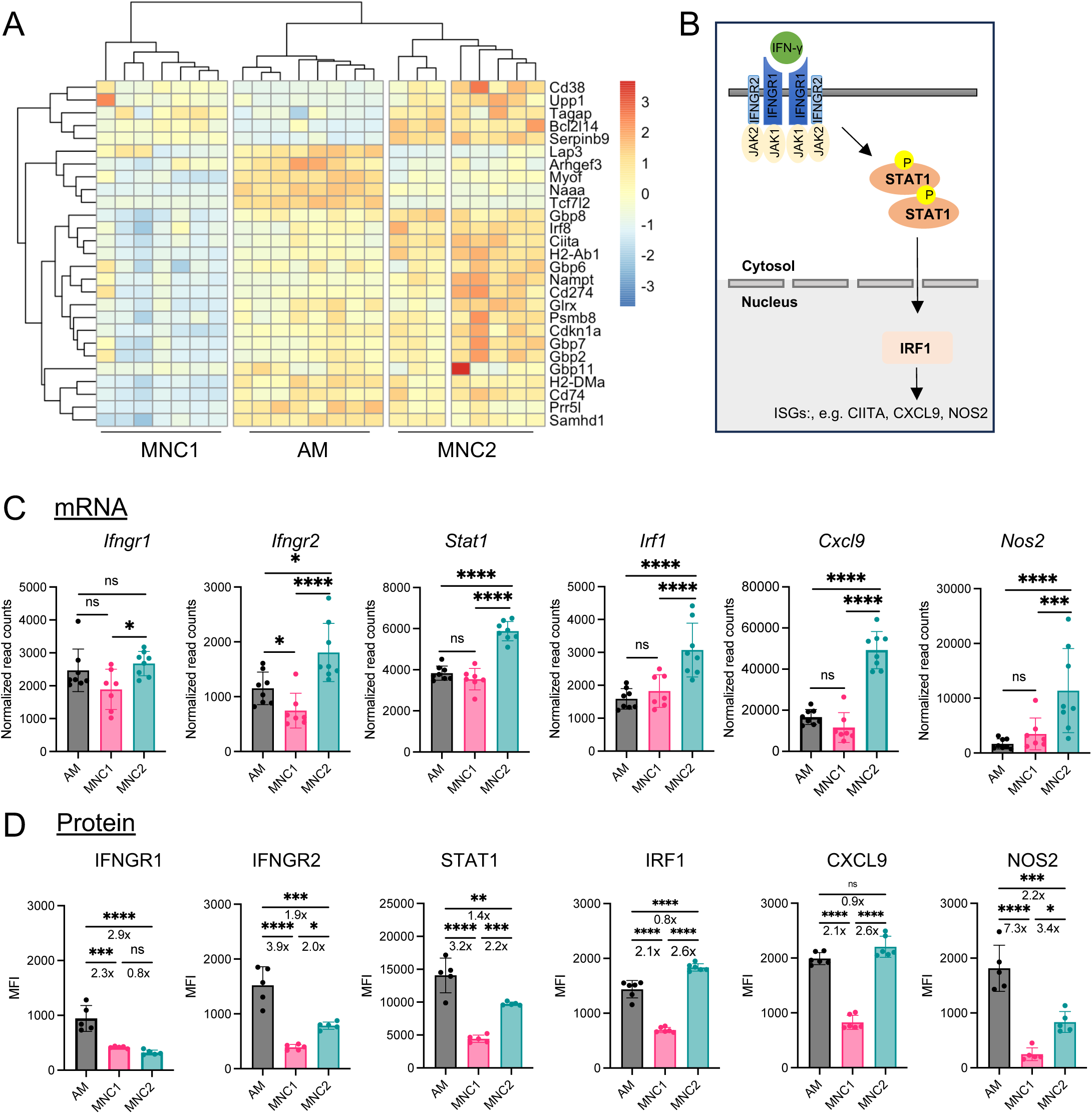
Key proteins in IFNγ signaling pathway are expressed at lower levels in MNC1. (A) Analysis of RNA-seq dataset GSE288567 revealed that IFNγ signature genes are expressed at lower levels in CD11c^lo^ monocyte-derived cells (MNC1) compared with alveolar macrophages (AM) and CD11c^hi^ MNC2 (28 dpi). 10,000 live cells from each subset were sorted from the lungs of C57BL/6 mice infected with Mtb H37Rv-mCherry (n = 7-8). (B) A schematic diagram shows the key signaling proteins regulating the response to IFNγ and representative IFNγ-responsive genes. (C) Messenger RNAs encoding key signaling proteins regulating the response to IFNγ and representative IFNγ-responsive genes are differentially expressed in AM, MNC1, and MNC2. Shown are mRNA normalized read counts for different cell subsets (28 dpi) (n=7-8). (D) Key signaling proteins regulating the response to IFNγ and protein products of representative IFNγ-responsive genes are differentially expressed in AM, MNC1, and MNC2. Flow cytometry was used to measure median fluorescent intensity (MFI) of proteins in different lung cell subsets from mice infected with H37Rv-ZsGreen. Fold change values were for AM vs. MNC1, MNC2 vs. MNC1, and AM vs. MNC2 (n=4-5). Results are presented as mean ± SD. *p<0.05, **p<0.01, ***p<0.0001, ****p<0.0001 by Wald test via DESeq2 for (C) and one-way ANOVA with Tukey’s multiple comparison test for (D); ns = not significant.

### MNC1 are deficient in responding to IFNγ ex vivo

Based on the above findings, we hypothesized that lung MNP subsets isolated from lungs during chronic Mtb infection either differ in their ability to respond to IFNγ, or that differential responses were due to differential exposure to IFNγ, perhaps due to their positioning or access to IFNγ in vivo. To distinguish these two possibilities, we compared responses to ex vivo IFNγ stimulation for lung MNP subsets by flow cytometric analysis of tyrosine phosphorylated STAT1 (pSTAT1), a key early signaling step in IFNγ cell activation. We prepared lung single-cell suspensions from Mtb-infected mice (28 dpi) and treated them with or without IFNγ (Figure 2A). Without IFNγ stimulation, AM and MNC2 showed higher pSTAT1 levels than MNC1, likely reflecting IFNγ stimulation in vivo. With ex vivo IFNγ stimulation, AM, MNC1, and MNC2 exhibited a 5.1-fold, 2.3-fold, and 2.5-fold increase of pSTAT1, respectively (Figure 2B-2C and Figure S2); however, MNC1 still showed a lower pSTAT1 signal than AM or MNC2 (Figure 2B). Interestingly, although infected AM and infected MNC2 responded to IFNγ better than their bystander counterparts, the subset difference in IFNγ responsiveness between AM, MNC1, and MNC2 did not depend on the presence of Mtb in individual cells (Figure S3). We also observed a similar phenotype at later time points of Mtb infection (Figure S4).

**Figure 2.**
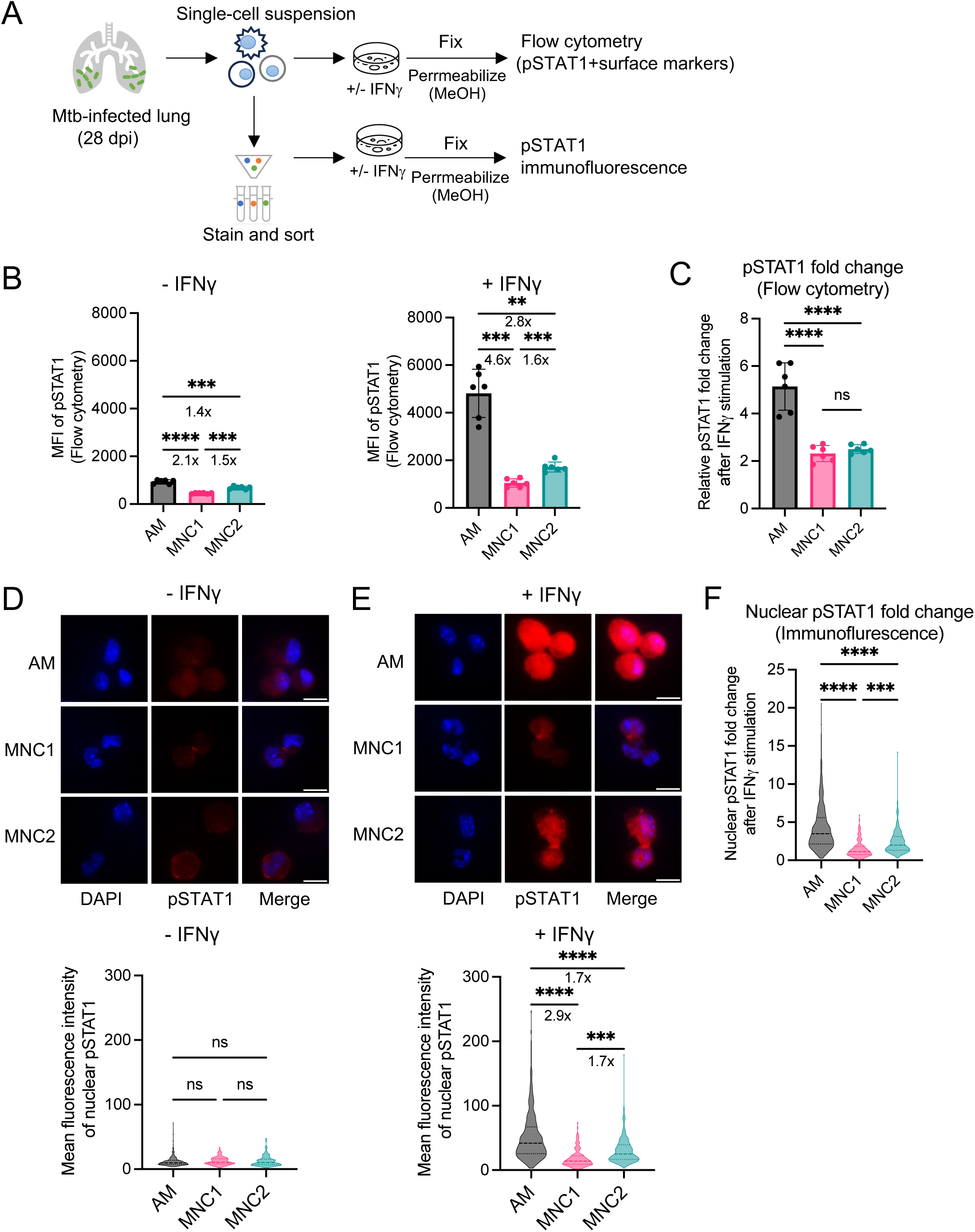
STAT1 phosphorylation is deficient in MNC1 treated with IFNγ ex vivo. (A) Workflow for detecting pSTAT1 by flow cytometry and immunofluorescence. C57BL/6 mice were infected with Mtb H37Rv-ZsGreen via aerosol for 28 days. Lung single-cell suspensions were incubated with 20 ng/mL IFNγ for 15 min and subjected to pSTAT1 by flow cytometry analysis or immunofluorescence. (B) MFI of pSTAT1 by flow cytometry for each lung cell subset in the absence or presence of IFNγ stimulation as described in panel A. Fold change values were for AM vs. MNC1, MNC2 vs. MNC1, and AM vs. MNC2 (n=6). (C) Fold change of pSTAT1 MFI level by flow cytometry after cells were stimulated with IFNγ ex vivo as described in panel B (n=6). (D) Representative images of nuclear pSTAT1 immunofluorescence staining for each cell subset sorted from Mtb-infected mice (28 dpi). Bottom panel shows the quantitation of mean fluorescence intensity of nuclear pSTAT1 from n>145 cells. (E) Representative images of nuclear pSTAT1 immunofluorescence staining for each cell subset sorted from Mtb-infected mice (28 dpi) and stimulated with 20 ng/mL IFNγ for 15 min. The bottom panel shows the quantitation of the mean fluorescence intensity of nuclear pSTAT1 from >155 cells. Fold change values were for AM vs. MNC1, MNC2 vs. MNC1, and AM vs. MNC2. (F) Fold change of nuclear pSTAT1 MFI by immunofluorescence staining after cells were stimulated with IFNγ ex vivo as described in panel E. Results are presented as mean ± SD. Brown-Forsythe and Welch ANOVA tests with Dunnett’s T3 multiple comparisons test (B, D-F) and One-way ANOVA with Tukey’s multiple comparison test (C) were used to determine statistical significance. ***p<0.001, ****p<0.0001 by one-way ANOVA; ns = not significant.

Since pSTAT1 functions in the nucleus, we used immunofluorescence to quantify nuclear pSTAT1 in sorted mononuclear cells with or without ex vivo IFNγ. In the absence of added IFNγ, the nuclear pSTAT1 level was similar in cells in all three subsets (Figure 2D). However, after IFNγ stimulation, we observed that MNC1 cells exhibited lower nuclear pSTAT1 levels than AM and MNC2, while MNC2 had an intermediate level between those of AM and MNC1 (Figure 2E). Furthermore, in response to IFNγ stimulation, MNC1 showed a lower fold change of nuclear pSTAT1 (1.5-fold) than AM (4.3-fold) or MNC2 (2.4-fold) (Figure 2F). Collectively, these results indicate that responsiveness to IFNγ is impaired at a proximal step of signal transduction in MNC1 compared with AM or MNC2, while MNC2 has an intermediate level of IFNγ responsiveness.

### MNC1 are impaired for activating Mtb antigen-specific CD4 T cells

One effect of IFNγ is to upregulate MHC-II, which mediates antigen presentation and CD4 T cell activation^19–21^. Therefore, we hypothesized that MNP subsets that differ in IFNγ responsiveness also differ in MHC-II expression and antigen presentation. GO analysis revealed that transcripts encoding multiple components of the MHC-II antigen presentation pathway are differentially regulated when comparing AM or MNC2 to MNC1 (Table S2). We found that class II transactivator (CIITA), the master transcription factor for MHC-II expression and other MHC-II antigen presentation-related genes is underexpressed in MNC1 (Figure 3A). Using flow cytometry, we confirmed that MHC-II and CD74 (the MHC-II invariant chain) are expressed at lower levels in MNC1 cells compared to AM and MNC2 (Figure 3B-3C and Figure S2). Differential MHC-II expression was also observed at 42 dpi and 56 dpi (Figure S5). Next, we sought to determine whether reduced MNC1 expression of MHC-II impairs antigen presentation to CD4 T cells. To specifically study antigen presentation independently of potential differences in antigen processing, we incubated sorted cells from MNP subsets from Mtb-infected mice with C7 or P25 epitope peptides (from ESAT-6 or Ag85B, respectively), and antigen-specific Th1-polarized C7 or P25 TCR transgenic CD4 T cells, then used IFNγ ELISA as a readout of T cell activation (Figure 3D). Consistent with the levels of expression of MHC-II pathway molecules, we observed lower responses by antigen-specific CD4 T cells when stimulated by MNC1 compared with AM or MNC2 (Figure 3E-3F), confirming that MNC1 cells have an impaired ability to present Mtb antigens to CD4 T cells.

**Figure 3.**
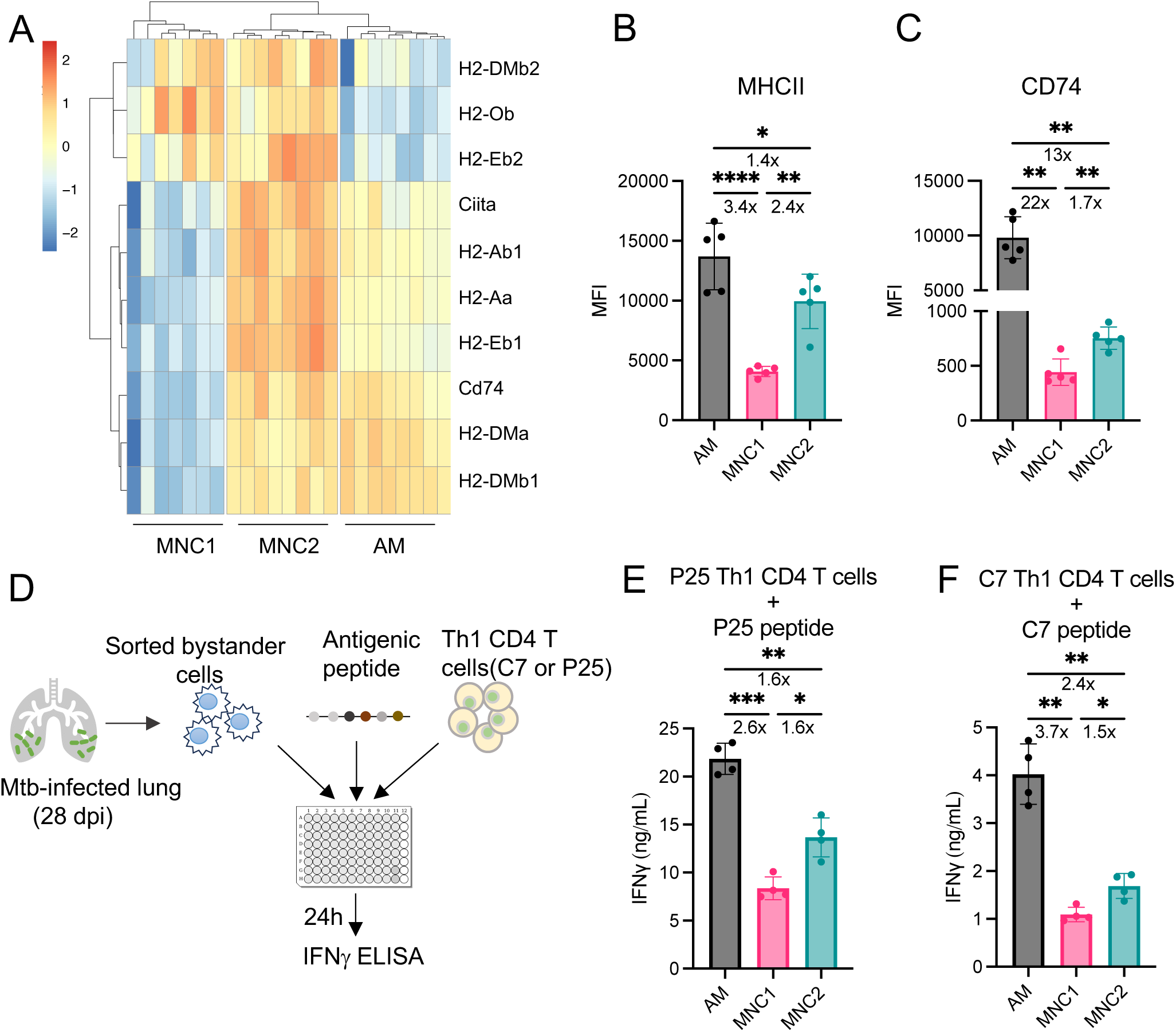
Impaired responsiveness to IFNγ in MNC1 associates with defective MHC-II antigen presentation. (A) Heatmap of the Z-scores from variance-stabilized read counts for genes related to MHC-II antigen presentation (adjusted p-value ≤ 0.05 and |log2 fold change|>0.5). (B-C) MNC1 express less surface MHC-II (B) and CD74 (C) than AM and MNC2. Flow cytometry was used to measure MFI of MHC-II and CD74 in each cell subset from mice infected with Mtb H37Rv-ZsGreen (28 dpi). Fold change values were for AM vs. MNC1, MNC2 vs. MNC1, and AM vs. MNC2 (n=5). (D) Workflow of antigen-specific T cells and lung mononuclear phagocyte co-culture. 1×10^4^ cells were sorted from mice infected with Mtb H37Rv-ZsGreen and incubated with 2×10^5^ of the indicated Th1 polarized TCR transgenic antigen-specific T cells and 10ug/mL antigenic peptide. After 24h, the supernatant was filtered and assayed for IFNγ using ELISA. (E-F) Impaired antigen presentation to CD4 T cells by MNC1. Supernatants were harvested from the co-culture of antigen-specific T cells and different sorted cell subsets, and IFNγ level was measured by ELISA. Fold change values were for AM vs. MNC1, MNC2 vs. MNC1, and AM vs. MNC2 (n=4). Results are presented as mean ± SD. One-way ANOVA with Tukey’s multiple comparison test (B) and Brown-Forsythe and Welch ANOVA tests with Dunnett’s T3 multiple comparisons test (C, E-F) were used to determine statistical significance. *p<0.05, **p<0.01, ****p<0.0001; ns = not significant.

### Type I IFN inhibits MHC-II expression in primary macrophages and lung MNPs

We next investigated the potential mechanisms regulating MHC-II expression in lung MNP subsets. From our bulk RNA-seq data, we found a higher type I IFN signature in MNC1 and MNC2 compared to AM (Figure 4A), consistent with results from our prior study^4^. Since type I IFN has been reported to cause down-regulation of IFNGR ^22–24^, we reasoned that type I IFN might suppress IFNγ-mediated MHC-II upregulation in MNPs/macrophages in Mtb infection. To test this, we differentiated bone marrow-derived macrophages (BMDM) in the presence of IFNγ and/or IFNβ, and quantified MHC-II expression by flow cytometry. As expected, IFNγ significantly increased MHC-II in wild-type (WT) BMDM (Figure 4B). IFN-β alone slightly reduced MHC-II expression and inhibited IFNγ-mediated MHC-II upregulation. Deletion of the type I IFN receptor, *Ifnar1*, completely negated the inhibitory effect of IFNβ on MHC-II expression in BMDM (Figure 4C).

**Figure 4.**
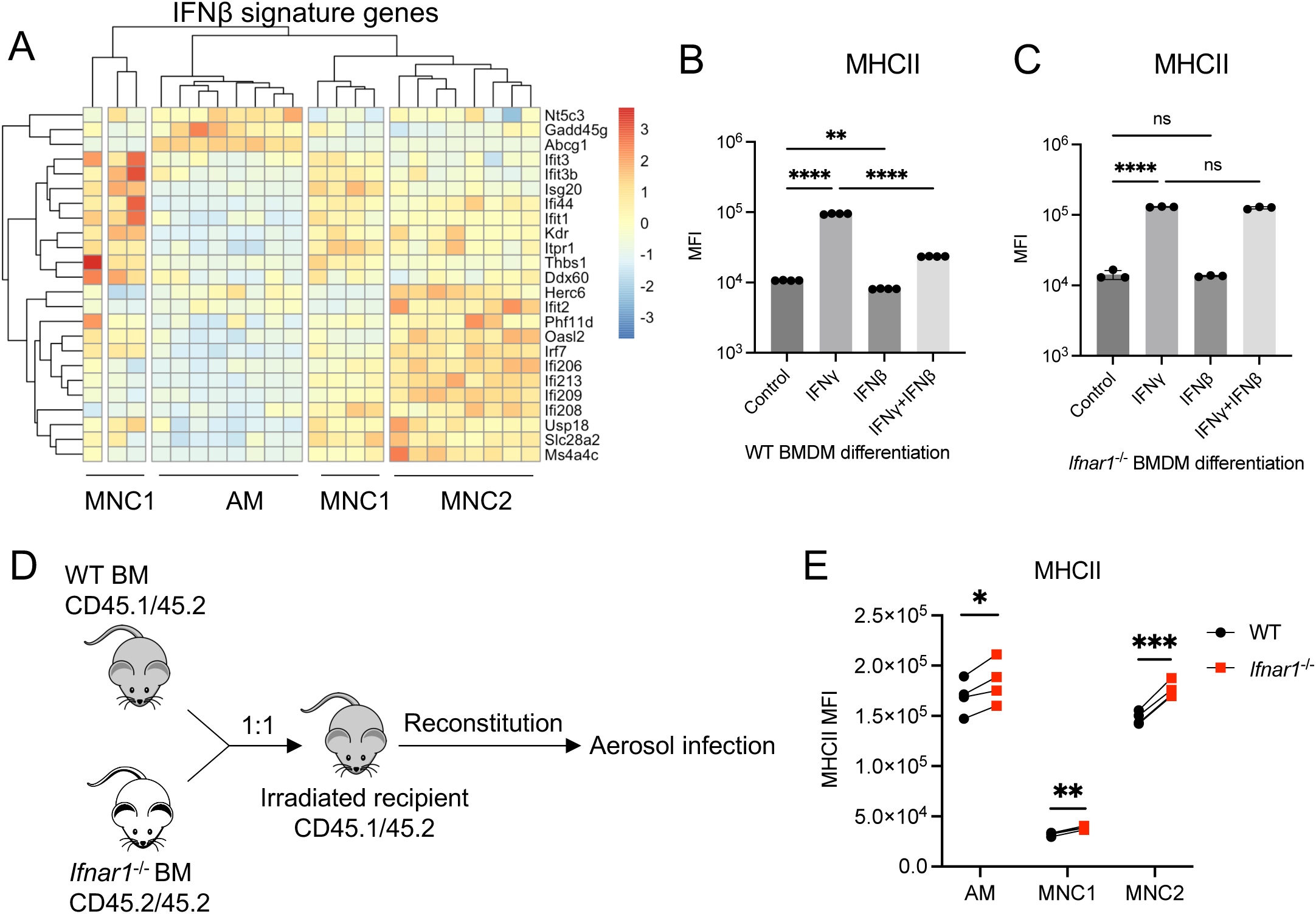
Type I IFN contributes to defective IFNγ responses and MHC-II expression in BMDM and MNP subsets. (A) Heatmap of the Z-scores from variance-stabilized read counts for IFNβ signature genes (adjusted p-value ≤ 0.05 and |log2 fold change|>0.5, n=7-8). (B-C) Type I IFN suppresses MHC-II expression induced by IFNγ in vitro. Wild-type (WT) or *Ifnar1*^−/-^ bone marrow cells were differentiated in bone marrow-derived macrophage (BMDM) media containing the indicated interferon at 50 pg/mL for 6-7 days. The expression levels of MHC-II in BMDM were determined by flow cytometry and expressed as median fluorescence intensity (MFI) (n=4). (D) Experimental workflow. CD45.1/45.2 mice were irradiated and reconstituted with 1:1 mixed bone marrow from WT CD45.1/45.2 mice and *Ifnar1*^−/-^ CD45.2/45.2 mice. After 6-week reconstitution, chimeras were aerosol challenged with Mtb H37Rv-ZsGreen. (E) Type I IFN regulates MHC-II expression in AM, MNC1, and MNC2 in vivo. Flow cytometry was used to measure MHC-II MFI in each subset from WT: *Ifnar1*^−/-^ mixed bone marrow chimeric mice infected with Mtb H37Rv-ZsGreen for 28 days (n=4). Results are presented as mean ± SD. *p<0.05, **p<0.01, ***p<0.0001, ****p<0.0001 by one-way ANOVA with Tukey’s multiple comparison test for (B-C) or by paired t-test for (E); ns = not significant.

To further investigate the effect of type I IFN on MHC-II expression on different MNP subsets in vivo, we generated mixed bone marrow chimeric mice (Figure 4D). Irradiated WT recipient mice received WT and *Ifnar1*^−/-^ mixed bone marrow cells. After reconstitution, the mice were infected with Mtb by aerosol and harvested 28 dpi. All *Ifnar1*^−/-^ lung MNP subsets showed higher MHC-II expression compared to the analogous WT subsets (Figure 4E and Figure S6). Together, these results indicate that type I IFN negatively regulates IFNγ-mediated MHC-II expression in BMDM in vitro and in lung MNP subsets in vivo in the context of Mtb infection.

### CoMtb-mediated prior immunity improves IFNγ responsiveness of MNC cells but not AM

Since monocytes and macrophages are functionally reprogrammable, we investigated whether pre-existing immunity alters IFNγ responsiveness of MNP subsets, using the contained Mtb (CoMtb) model^25^. This model is established by intradermal infection of Mtb in the mouse ear, which results in Mtb trafficking to the local draining lymph nodes without bacterial dissemination to other organs, including the lungs.

Since MNC cells are derived from monocytes^4,8,26,27^, we first asked whether CoMtb regulates IFNγ responsiveness in bone marrow monocytes ex vivo. For this, we established CoMtb for 4-6 weeks (Figure 5A), then harvested bone marrow cells from CoMtb and control mice. Without IFNγ stimulation, we observed higher pSTAT1 in CoMtb bone marrow monocytes compared with control monocytes (Figure 5B). After IFNγ stimulation, CoMtb monocytes showed greater STAT1 activation than control monocytes; BMDM cultured from CoMtb mice also exhibited greater pSTAT1 than did BMDM from control mice. These data suggest that CoMtb conditions monocytes and macrophages to improve their IFNγ responsiveness in vivo (Figure 5B-5C).

**Figure 5.**
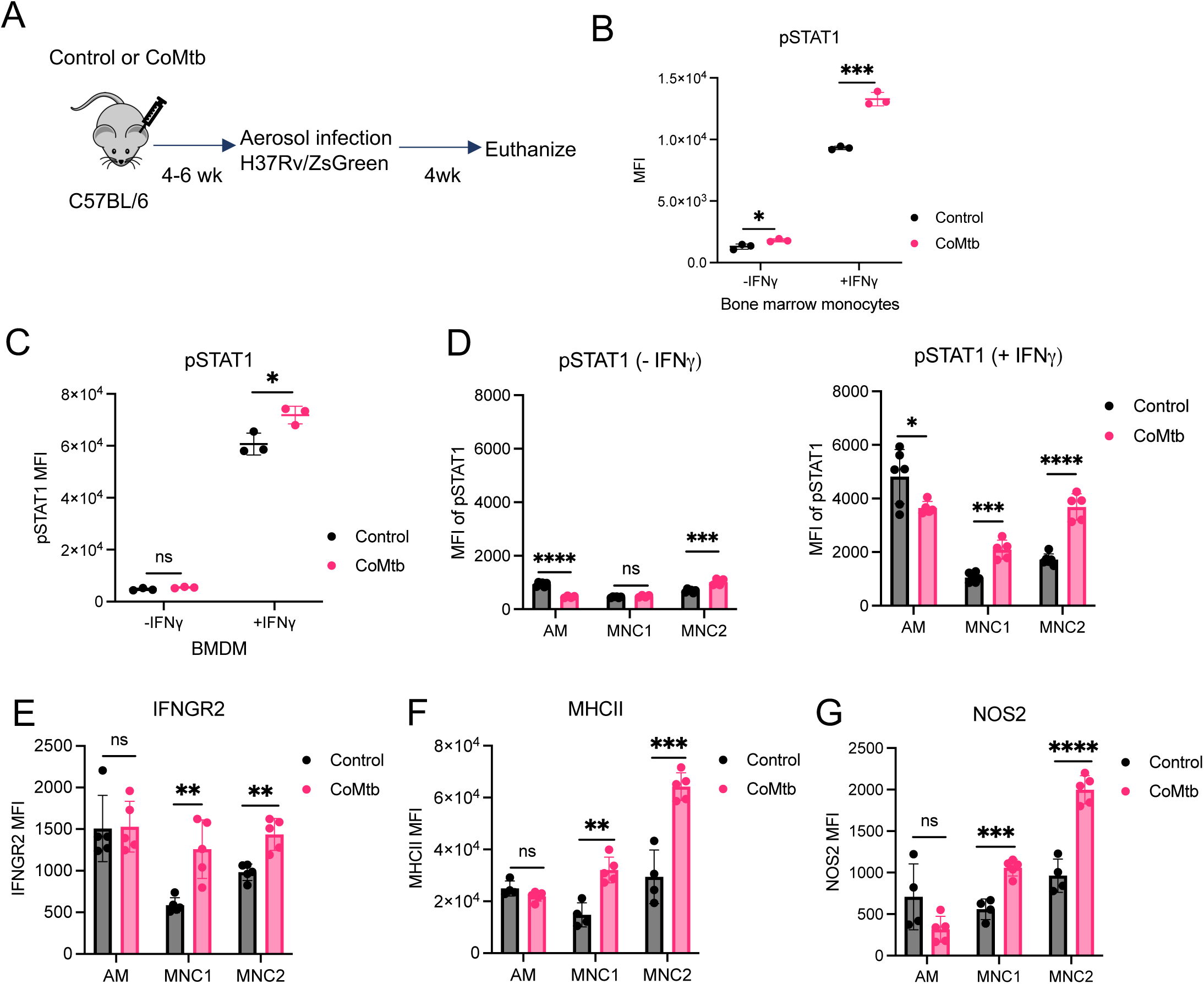
CoMtb improves IFNγ responsiveness of MNC but not AM. (A) For the contained Mtb (CoMtb) studies, mice were inoculated intradermally in the ear with 10,000 CFU of Mtb H37Rv. After 4-6 weeks, control and CoMtb mice were infected with Mtb H37Rv-ZsGreen by aerosol and euthanized at 28 dpi. (B) CoMtb increases IFNγ responsiveness in bone marrow monocytes. Bone marrow cells from control or coMtb mice were stimulated with or without 20 ng/mL IFNγ for 15 min, followed by flow cytometric measurement of pSTAT1 MFI (n=3). (C) CoMtb increases IFNγ responsiveness in BMDM. Bone marrow cells from control or coMtb mice were differentiated into BMDM for 6 days. Then BMDM were harvested and stimulated with or without 20 ng/mL IFNγ for 15 min, followed by flow cytometric measurement of pSTAT1 MFI (n=3). (D) CoMtb improves IFNγ responsiveness in MNC1 and MNC2. Lung samples from control or coMtb mice infected with H37Rv-ZsGreen by aerosol (28 dpi) were stimulated with or without 20 ng/mL IFNγ for 15 min, followed by flow cytometric measurement of pSTAT1 MFI (n=5-6). (E) CoMtb increases expression of IFNGR2 in MNC1 and MNC2. Shown are IFNGR2 MFI by flow cytometry for each lung cell subset from control or coMtb mice infected with H37Rv-ZsGreen by aerosol (28 dpi) (n=5-6). (F) CoMtb enhances MHC-II expression in MNC1 and MNC2. MHC-II MFI was measured by flow cytometry (28 dpi) (n=4-5). (G) MNC1 and MNC2 from CoMtb mice show a higher expression of NOS2 by flow cytometry (28 dpi) (n=4-5). Results are presented as mean ± SD. *p<0.05, **p<0.01, ***p<0.0001, ****p<0.0001 by unpaired t tests; ns = not significant.

We further investigated the effect of CoMtb on IFNγ responsiveness in lung MNP subsets from mice after subsequent aerosol Mtb infection. Using flow cytometry detection of pSTAT1 to quantitate IFNγ responses, we found that CoMtb improved responses to IFNγ in MNC1 and MNC2 cells, but not in AM, (Figure 5D). CoMtb also increased expression of IFNGR2 and the IFNγ-regulated proteins MHC-II and NOS2 (Figure 5B). These results indicate that CoMtb improves IFNγ responsiveness in monocyte-derived cells that traffic to the lungs after aerosol Mtb challenge.

### CoMtb-mediated control of Mtb in lung monocyte-derived cells requires IFNγ signaling

As indicated by whole lung CFU, we confirmed that CoMtb reduced Mtb burden in the lung after aerosol infection (Figure 6A); we also found that CoMtb reduced the number of total Mtb-infected cells in the lungs and lower bacterial fluorescence intensity (indicative of the per-cell bacterial burden) in the infected cells (Figure 6B-6C). We further found that CoMtb decreased the number of infected AM, MNC1, MNC2, and neutrophils compared to controls (Figure 6D). In addition, the infection frequency (i.e., the percentage of cells in a given subset that contain Mtb) was reduced in all subsets when comparing CoMtb mice to control mice (Figure 6E). Moreover, we found that CoMtb reduced Mtb burdens in AM (1.2-fold decrease), neutrophils (1.5-fold decrease), and especially in MNC1 (2.7-fold decrease) and MNC2 (1.8-fold decrease) (Figure 6F). In line with these results, chimeric mice that received bone marrow from CoMtb mice showed lower lung CFU burdens than mice that received control bone marrow cells (Figure 6G-6H). Thus, these findings indicate that improved IFNγ responsiveness can enhance Mtb control in monocyte-derived lung cells.

**Figure 6.**
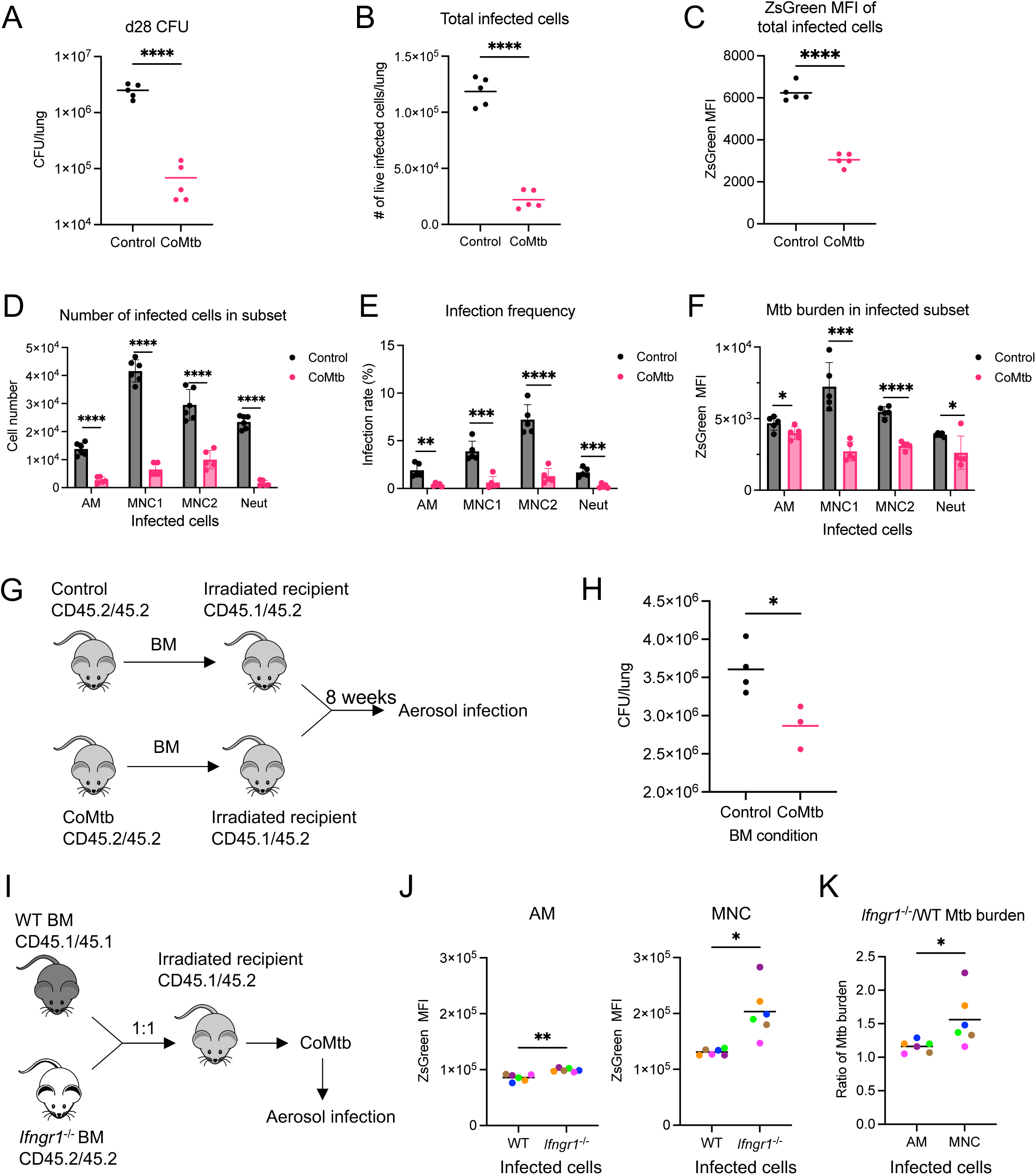
CoMtb-mediated prior immunity improves Mtb control in lung myeloid cells. (A) Total lung CFU for control or CoMtb mice infected with Mtb H37Rv-ZsGreen via aerosol for 28 days (n=5). (B) The number of total live infected cells from the same lung in panel A (n=5). (C) Bacterial burden indicated by ZsGreen MFI by flow cytometry for total infected cells in panel B (n=5). (D) Number of infected cells in each subset from the lungs of control and CoMtb mice infected with Mtb H37Rv-ZsGreen via aerosol for 28 days (n=5-6). (E) Frequency of Mtb-infected cells in each lung subset from control and CoMtb mice infected with Mtb H37Rv-ZsGreen via aerosol for 28 days (n=5). (F) Bacterial burden in each infected lung cell subset from control and CoMtb mice infected with Mtb H37Rv-ZsGreen via aerosol for 28 days (n=5). (G) Irradiated mice received 5 × 10^6^ T cell-depleted bone marrow cells from control or coMtb mice. After reconstitution for 8 weeks, mice were infected with aerosol Mtb for 28 days. (H) Lung Mtb burden for Mtb-infected chimeric mice that received bone marrow cells from control or coMtb mice (28 dpi) (n=3-4). (I) To generate mixed bone marrow chimeric mice, recipient (CD45.1/45.1) mice were irradiated and reconstituted with 1:1 mixed bone marrow from WT mice and *Ifngr1*^−/-^ CD45.2/45.2 mice for 8 weeks. After 4 weeks of CoMtb, mice were aerosol challenged with Mtb H37Rv-ZsGreen. (J) Bacterial burden indicated by ZsGreen MFI in infected AM and MNC cells from infected bone marrow chimeric mice. There were insufficient infected MNC1 in CoMtb mice for separate analyses, so MNC1 and MNC2 were combined as MNC in these studies (n=6). (K) Ratio of *Ifngr1*^−/-^/WT Mtb burden in different Mtb-infected subsets (ZsGreen^+^) from infected bone marrow chimeric mice (n=6). Results are presented as mean ± SD. *p<0.05, **p<0.01, ***p<0.0001, ****p<0.0001 by unpaired t-test (A-F, H), or by paired t-test for (J-K); ns = not significant.

We next sought to determine whether IFNγ responsiveness is required for CoMtb-mediated Mtb control in MNC cells. For this, we generated mixed bone marrow chimeric mice (Figure 6I). Irradiated WT mice received WT and *Ifngr1*^−/-^ mixed bone marrow cells (1:1). After reconstitution and CoMtb establishment, they were infected with Mtb by aerosol and harvested at 28 dpi. This revealed higher bacterial burdens in *Ifngr1*^−/-^ myeloid cells compared to WT myeloid cells from the same chimeric mouse (Figure 6J and Figure S7). Similarly, we observed a higher Mtb burden in *Ifngr1*^−/-^ MNC cells compared to WT counterparts (Figure 6J-6K). However, we observe a slight difference in Mtb burden between WT and *Ifngr1*^−/-^ AM at 28 dpi (Figure 6J), consistent with our data showing that CoMtb plays a minimal role in AM at the later stage of Mtb infection, in contrast to the effect of CoMtb on AM at an early stage of infection^25^. Collectively, these results suggest that CoMtb-mediated prior immunity enhances Mtb control in MNC cells, which depends on IFNγ signaling.

## Discussion

A predominant adaptive immune response to Mtb infection in mice, nonhuman primates, and humans is represented by T lymphocytes that produce IFNγ^15,19,28–34^, a cytokine that plays an essential role in activating macrophages to restrict diverse intracellular pathogens. We and others have determined that Mtb resides in multiple macrophage subsets in the lungs^4,7–10,20,35–38^. However, immunity to TB is often incomplete despite abundant IFNγ production, and increasing IFNγ levels alone does not necessarily improve bacterial control in the lungs ^1 17^ . Together, these results suggest that while IFNγ is necessary, its efficacy may be limited by impaired cellular responses and other inhibitory mechanisms in vivo that blunt the effects of IFNγ. This paradox highlights the need to better understand the mechanisms regulating IFNγ responsiveness in lung macrophage subsets to develop more effective TB therapies and vaccines.

Here, we identified a previously undefined mechanism that contributes to the suboptimal efficacy of IFNγ in immunity to Mtb. Using an unbiased initial approach, bulk RNA sequencing of flow-sorted mononuclear-macrophage cell subsets (AM and two subsets of lung monocyte-derived cells), we detected evidence that one of the monocyte-derived cell subsets (termed MNC1, formerly termed recruited macrophages) expresses lower levels of transcripts for IFNγ-responsive genes when compared with AM or the other monocyte-derived cell subset (MNC2). Since this could have reflected differential exposure of these cells to IFNγ in vivo, we treated lung cell suspensions ex vivo with IFNγ and found that MNC1 cells are less responsive to IFNγ as manifested by reduced tyrosine phosphorylation of STAT1. We also found that at the levels of mRNA and cell surface expression, MNC1 cells express lower levels of the IFNγ receptor subunit, IFNGR2, and other key signaling molecules in the response to IFNγ, including STAT1 and IRF1. These results indicate that in MNC1 cells, defective responses to IFNγ are associated with reduced expression of the proximal signaling machinery required for cell responses to IFNγ, and this results in reduced responses to IFNγ in these cells. Our findings complement in vitro observations that cultured macrophages infected with Mtb are defective in responding to IFNγ through multiple mechanisms. These mechanisms include prolonged TLR2 signaling initiated by Mtb lipoproteins and other cell wall components^39–44^, and inhibition of macrophage responses to IFNγ by IL-6 ^45^. Notably, these mechanisms inhibit IFNγ-responsive gene expression at one or more steps downstream of STAT1 activation and function^46,47^. Although the in vivo relevance of these mechanisms inhibiting macrophage responses to IFNγ has not been established, our study highlights a distinct, subset-specific pathway of IFNγ signaling impairment in MNC1 cells during chronic Mtb infection (Figure 7).

**Figure 7.**
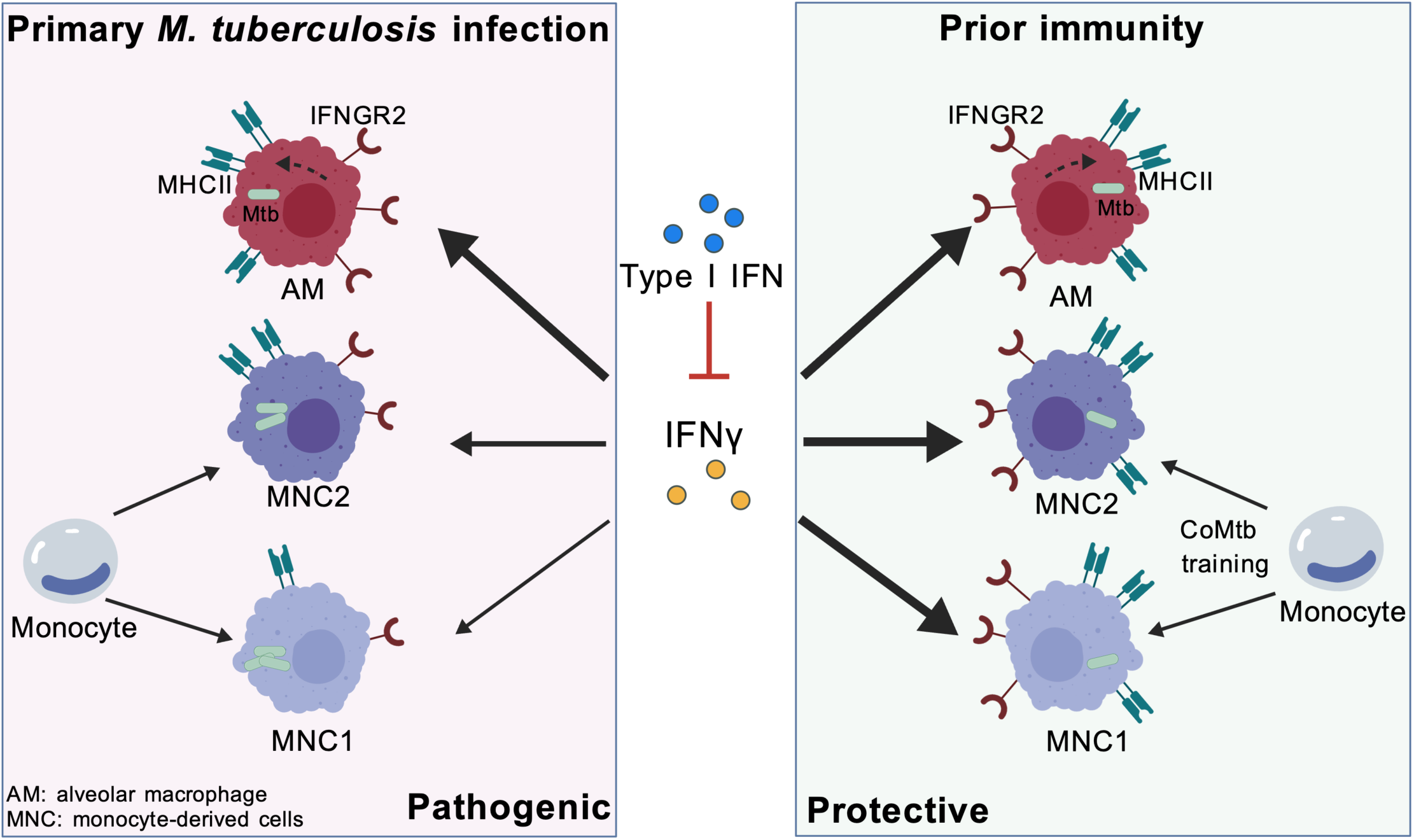
Proposed model of subset-specific IFNγ responsiveness in Mtb infection. In Primary infection, MNC1 exhibit low levels of IFNγ signaling components (e.g., IFNGR2). This impaired responsiveness, compounded by Type I IFN-mediated suppression of MHC-II expression, renders MNC1 cells highly permissive to intracellular Mtb growth and reduces their ability to be activated by CD4 T cells. In contrast, MNC2 and AM maintain higher IFNγ responsiveness. In CoMtb model, monocyte-derived cells undergo training, showing enhanced IFNγ responsiveness and upregulated MHC-II expression. This shift results in a more robust activated phenotype, leading to improved restriction of Mtb replication and superior protective immunity. Created with biogdp.com.

Recent studies reveal that the Mtb-killing ability in distinct macrophage subsets differs with the stage of infection. In early infection prior to development of adaptive T cell responses, AM are permissive to Mtb^35–37^, while after adaptive T lymphocytes are primed and recruited to the lungs, AM restrict Mtb more effectively than do monocyte-derived cells ^4,7–10^. Specifically, we previously reported that MNC1 cells harbor more viable Mtb per infected cell than do AM and MNC2 cells during chronic infection^8^. Adaptive T cells contribute to determining the fate and state of Mtb in distinct lung mononuclear cell subsets, and T cells have a greater impact on suppressing Mtb in AM than in recruited mononuclear cells in the lungs^7^. Our work extends these findings by demonstrating that subset-specific differences in IFNγ responsiveness may explain why T cell-mediated adaptive immunity has a greater impact on bacterial control in AM than in monocyte-derived cells, indicating that impaired IFNγ responsiveness contributes to Mtb permissiveness.

Although we have not yet fully identified the regulatory pathway(s) responsible for reduced expression of the signaling molecules required for responses to IFNγ, we found that type I IFN signaling contributes to reduced IFNγ-mediated MHC-II expression in MNC1 cells, impairing their ability to present antigens to antigen-specific CD4+ T cells. This provides a mechanistic link between the known detrimental role of Type I IFN in TB and the failure of CD4^+^ T-cell-mediated control at the cellular level. Although MNC2 cells express higher mRNA levels of IFNγ pathway genes, they are less responsive to IFNγ stimulation compared to AM. This suggests that MNC2 may be exposed to higher levels of IFNγ than AM cells, likely due to their localization. However, MNC2 show a greater type I IFN signaling than AM, which may antagonize IFNγ signaling. Interestingly, IFNγ responsiveness in MNC1 is not fully restored by deletion of *Ifnar1*, suggesting that other mechanisms may contribute to regulating IFNγ responsiveness in lung MNC subsets in vivo during Mtb infection.

An important discovery from this study is that the impaired IFNγ responsiveness of MNC1 cells is not an irreversible differentiation trait but rather a malleable phenotype that can be reprogrammed by prior immunity (Figure 7). We found that the responsiveness of MNC1 cells to IFNγ can be modulated: pre-existing immunity established by CoMtb modifies bone marrow precursors of MNC1 and MNC2 cells to improve their responses to IFNγ and improve their ability to control Mtb. In this manner, our results are consistent with trained immunity^48^, since we found that the impact of CoMtb on myeloid cells influenced ex vivo responses to IFNγ and that transfer of bone marrow cells from mice with CoMtb improved control of Mtb after aerosol challenge. While deleting the IFNγ receptor slightly increased bacterial burden in AM from coMtb mice, the impact of IFNγR deletion on bacterial control is more pronounced in monocyte-derived cells. This aligns with a recent study showing that CoMtb induces IFNγ signaling in MNC cells from Mtb-susceptible mice^49^. These findings indicate that the mechanisms of enhanced immunity in CoMtb mice are cell-type specific. MNC1 cells face a critical ‘bottleneck’ due to impaired IFNγ signaling during primary infection; thus, restoring this responsiveness via CoMtb is a primary driver of their improved control. In contrast, while IFNγR signaling contributes to AM function at early stages^25,50,51^, it plays a more limited role in the CoMtb-mediated protection of AMs during the later chronic stage of infection. However, we cannot exclude the possibility that CoMtb may alter other cell-intrinsic mechanisms, such as lysosome biogenesis and autophagy, to synergize with improved IFNγ responsiveness and enhance bacterial control, which warrants further study. Together, these results indicate that modulation of MNC1 cells can be achieved, suggesting that this may be an especially valuable target for host-directed TB therapies.

We also note that the cell subset (MNC1) that is deficient in responses to IFNγ is the same subset that we previously reported as being deficient in lysosome biogenesis. Since there are no known regulatory connections between IFNγ responsiveness and lysosome biogenesis, our results are most consistent with a model in which both pathways are regulated by a common developmental pathway during differentiation of bone marrow monocytes into monocyte-derived macrophages in the lungs of Mtb-infected mice. Further investigation of the differentiation or maturation of monocytes recruited to the lungs in response to Mtb will be needed for a more thorough understanding of the determinants of cell responsiveness to IFNγ and of lysosome biogenesis.

A shortcoming of the studies reported here is that they only involved studies in mice. Recent single-cell sequencing studies have revealed heterogeneity of lung monocyte-derived macrophages in human TB^52–54^, and a potential difference in IFNγ responses among macrophage subsets in TB granulomas^52^. However, functional studies are required to characterize IFNγ responsiveness in live macrophage subsets isolated from resected lung tissues of human TB patients. These are limited by the rarity of appropriate human samples in proximity to facilities for the procedures required, and considerable dedicated effort will be required to determine which of our findings are applicable to humans. Nevertheless, our results provide further mechanistic insight into the potential functional significance of macrophage heterogeneity in the lungs in the context of Mtb infection and should drive further investigation of methods to optimally develop and test host-directed therapies for TB.

## Methods

### Mice

8-12-week-old C57BL/6 (Strain #:000664), *Ifnar1*^−/-^ (Strain #:028288) and *Ifngr1*^−/-^ (Strain #:003288) mice were obtained from Jackson Laboratory. Both sexes were used in this study. Extensive experiments and projects in our lab demonstrate no sex-specific differences in our murine TB models. Therefore, sex was not a consideration in this study; sample sex information has not been collected. Mice infected with Mtb were housed in the Animal Biosafety Level 3 facility. All animal experiments were approved by the Institutional Animal Care and Use Committee of the University of California, San Francisco.

### Bacterial strains, growth, and aerosol infection

Mtb H37Rv was transformed with pMV261::ZsGreen and cultured in Middlebrook 7H9 medium (BD) supplemented with 10% (v/v) ADC (albumin, dextrose, catalase), 0.05 % Tween 80, 0.2 % glycerol, and 50 μg/ml kanamycin^8^. Mice were infected with H37Rv or pMV261::ZsGreen via aerosol using a Glas-Col inhalation exposure unit, as previously described^4,8,26,38^. Mid-log cultures were centrifuged at 800g for 8 min. Clump-free cultures were transferred and diluted, and 5 mL of inoculum was added to the nebulizer. The target dose was 100 CFU/mouse. Infection dose was determined by plating lung homogenates 24 hours post-infection on 7H11 agar plates. CFUs were counted after incubation of plates at 37°C for 3 weeks.

### Lung homogenate preparation, flow cytometry, and cell sorting

Lung homogenates were prepared as described previously^4,8^. Lungs were perfused with 10 mL of PBS/2 mM EDTA via the right ventricle immediately after euthanasia. Lungs were processed with a gentleMACS dissociator (Miltenyi, lung program 1), and digested in 4 mL of RPMI-1640/5% HI-FBS containing 1 mg/mL collagenase D (Sigma-Aldrich) and 50 μg/mL DNase I (Sigma-Aldrich) for 30 min at 37 °C. Digested lung tissues were further dissociated with the gentleMACS dissociator (lung program 2) and then passed through a 70-μm cell strainer. Red blood cells were lysed with 3 mL ACK lysis buffer (Gibco) for 3 min and washed twice with RPMI-1640/5% HI-FBS.

For staining of surface antigens, cells were washed with PBS, then stained with 1:200 Zombie Aqua Fixable Viability Dye (BioLegend, 423101) and 1:100 anti-CD16/32 (clone 2.4G2; Fc receptor blockade) for 15 min at 4°C. After washing with PBS, cells were stained with flow antibodies from Table S3 in 1:1 PBS/Brilliant Stain Buffer (BD, 566349) for 30 min at 4°C. Samples were washed twice with PBS, then fixed overnight in 1% paraformaldehyde (PFA)/PBS. For cell sorting, cells were washed and resuspended in FACS buffer (PBS, 2% HI-FBS (v/v), 2 mM EDTA), and sorting was done using the BSL3 Sony MAC900 cell sorter.

For intracellular staining, cells were fixed and permeabilized using BD Fixation/Permeabilization Kit for 20 min at 4°C after surface staining, then washed and incubated with antibodies diluted in 1x BD Perm Wash/0.5% FBS for 30 min at RT. For IRF1, 0.1% Triton X-100/PBS was used for permeabilization. Samples were washed and acquired using an Aurora spectral flow cytometer (Cytek Biosciences) or Sony MA900.

### Ex vivo IFNγ stimulation for pSTAT1 flow cytometry

10^6^ cells per lung were stained with Zombie Aqua Fixable Viability Dye as described above at 4°C for 15 min. Samples were washed once with RPMI-1640/5% HI-FBS, then stimulated with 100 μL of 20ng/mL IFNγ (final concentration) (Biolegend, 575304) in media at 37°C for 15 min. Untreated samples were kept at 4°C. Samples were then mixed with 100 μL of 4% PFA/PBS and fixed at room temperature (RT) for 15 min. Cells were washed once with PBS, followed by permeabilization with 180 μL of cold methanol at 4°C for 12 min. After this, samples were washed twice and incubated with 50 μL of 1:50 anti-CD16/32 in PBS for 5 min, followed by the addition of antibodies, including AF647-pSTAT1 (Cell Signaling Technology, 8009S), diluted in 50 μL of Brilliant Stain Buffer and incubated for 30 min at RT. After washing twice with PBS, samples were resuspended in 1% PFA/PBS.

### Immunofluorescence

Lung cell subsets were sorted from mice infected with H37Rv-ZsGreen. 40,000 cells/well were seeded into 8-well Nunc Lab-Tek chamber slides (Thermo Scientific, 177445), and rested in a cell incubator for 1 h. Media were replaced with 100 μL of RPMI-1640/5% HI-FBS with or without 20ng/mL IFNγ, and cells were incubated for 15min at 37°C. Immunofluorescence was performed as previously described with slight modifications^8,55,56^. Media were removed and cells were fixed immediately with 4% PFA for 30min. Slides were immersed in 1% PFA/PBS and transferred to BSL2. Cells were washed twice with 200 μL PBS, followed by permeabilization with 200 μL of ice-cold methanol for 10 min at 4°C, and then washed 3 times with PBS. Samples were blocked with 5% goat serum/PBS for 30 min at RT, then incubated with 100 μL of pStat1 (Tyr701) Rabbit mAb (Cell Signaling Technology, 9167S, 1:200 diluted in 5% goat serum/PBS) for 2 h at RT. Then cells were washed three times with PBS and stained with 1:1000 Goat anti-Rabbit IgG AF647 (Invitrogen, A-21244) for 1 h at RT. Samples were washed three times with PBS and mounted with ProLong Diamond Antifade Mountant with DAPI.

### T cell and MNP coculture

T cell and MNP coculture were done as described previously^21,57^. ZsGreen^−^ Lung MNP subsets (AM, MNC1 and MNC2) were sorted from mice infected with H37Rv-ZsGreen. 10,000/well of MNP were cultured with P25TCR-Tg or C7TCR-Tg Th1 effector cells (200,000/well) in the presence of 10 μg/mL Ag85B peptide 25. Supernatants were collected at day 4 and assayed for IFNγ by ELISA.

### BMDM differentiation

BMDM were differentiated as described previously^8^. Briefly, bone marrow cells were cultured in BMDM media [DMEM (Gibco, 11965092), 10% heat-inactivated FBS (HI-FBS), and 20 ng/ml recombinant murine M-CSF (PeproTech)] for 6-7 days, and media change was done at day 3. For differentiation with interferon, BMDM media containing 50 pg/mL of IFNγ (Biolegend, 575304) or IFN-β (Sino Biological, 50708-MCCH) were used. At day 6-7, cells were washed twice with PBS and incubated with cold PBS/2mM EDTA for 5 min, then harvested by pipetting.

### Generation of bone marrow chimeras

Bone marrow chimeras were generated as described previously^14^. Bone marrow cells were harvested from the femurs and tibias of donor mice. After ACK lysis, cells were washed once with DMEM/10% HI-FBS, once with PBS, then filtered with a 70 μm strainer. Recipient mice were irradiated with a Precision X-Rad320 X-Ray irradiator using a dose of 10 Gy. Within 24h post-irradiation, 6×10^6^ total bone marrow cells were retro-orbitally injected into recipient mice using a 28G insulin syringe. For the chimera experiment using bone marrow cells from control or CoMtb mice, recipient mice received 5×10^6^ T cell-depleted bone marrow cells. Irradiated mice were given TMP-SMX antibiotic (Sulfamethoxazole and Trimethoprim Oral Suspension, 200 mg/40 mg per 5 mL, use 7.5 mL per 293 mL drinking water) for 3 weeks. Mice were allowed to reconstitute for at least 6-8 weeks.

### Establishment of CoMtb

CoMtb was established as described previously^25^. Briefly, mice were anesthetized with isoflurane. 1×10^4^ CFU of Mtb H37Rv were injected intradermally into the base of the ear in 10 μL PBS using a 31 G insulin syringe. After 4-6 weeks of establishment, mice were used for experiments.

### RNA sequencing and data analysis

Analysis of the RNA-sequencing dataset GSE220147 was performed as described previously^8^.

### Quantification and statistical analysis

Results are expressed as mean and standard error. Quantification of MFI from immunofluorescence images was done using ImageJ, and the data are background-subtracted. MFI data for the lung subsets are background-subtracted using the fluorescence minus one (FMO) control or uninfected control. Statistical analysis and p-values are stated in the figure legends and indicated in the figures, respectively. P <0.05 is considered significant.

## Data availability

The RNA-seq data supporting the findings of this study are available in the Gene Expression Omnibus (GEO) under the accession number: GSE220147. Source data are provided with this paper.

## Acknowledgments

We thank Lucas Chen, Michael Kwon, and Kelley Martinez for technical assistance. This work was supported by NIH grants R01 AI051242 (J.D.E.), U01 AI166309 (J.D.E.), P30 AI168440/R25 AI147375 (W.Z. and A.M.), T32 HL007185 (A.M.), F32 HL162424 (A.M.), and F31 AI172360

(Z.P.H.). The funders had no role in study design, data collection, interpretation, or the decision to submit the work for publication.

## Author contributions

Conceptualization, W.Z. and J.D.E.; methodology, W.Z., J.D.L., and Z.P.H.; investigation, W.Z., J.L., Z.P.H., and A.M.; resources: E.T.; writing—original draft, W.Z. and J.D.E.; writing—review & editing, W.Z. and J.D.E.; funding acquisition, W.Z., Z.P.H., A.M., and J.D.E.; supervision, W.Z. and J.D.E.

## Declaration of interests

The authors declare no competing interests.

## Notes

### Competing Interest Statement

The authors have declared no competing interest.

### Summary of Updates

Major adjustments in this work include: (1) modifying the Introduction and Discussion to delineate our novel contributions explicitly; (2) updating Figures 1-3 and adding a schematic diagram of the proposed model in Figure 7; (3) adding results of CoMtb chimera experiment to Figure 6 and representative histogram plots in Figure S2; (4) updating Figure legends and Method section.

