## Supplementary information for "Impaired IFNγ responsiveness of monocyte-derived lung cells limits immunity to *Mycobacterium tuberculosis*"

### Supplemental information

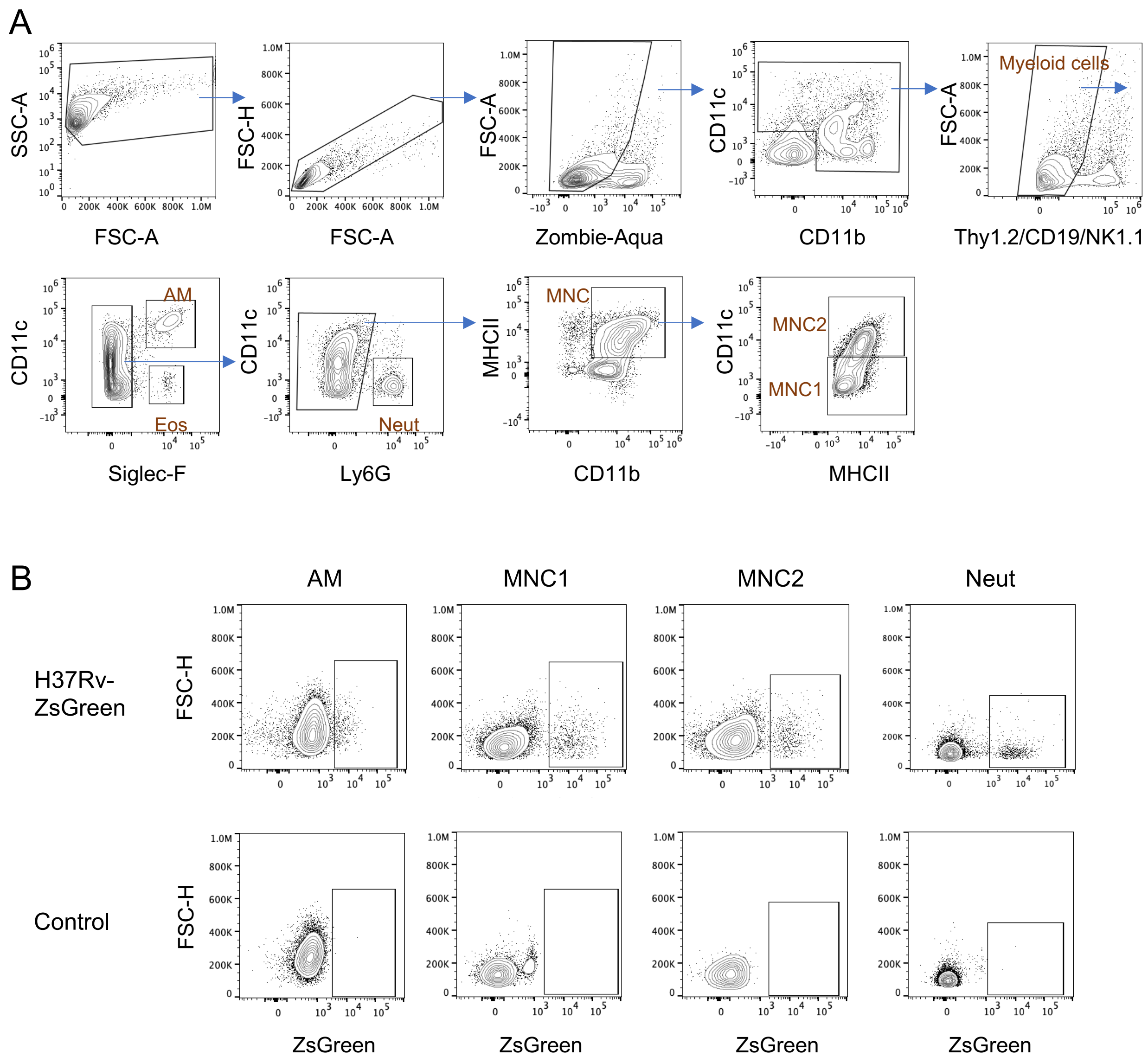


**Figure S1. Flow gating strategy for lung cell populations.**
(A) Representative flow cytometry plots to identify alveolar macrophages (AM), monocyte-derived cells (MNC1, MNC2), and neutrophils (Neut) from Mtb-infected mouse lungs.
(B) Representative plots of infected cells for each MNP subset from mice infected with H37Rv-ZsGreen (28 dpi).

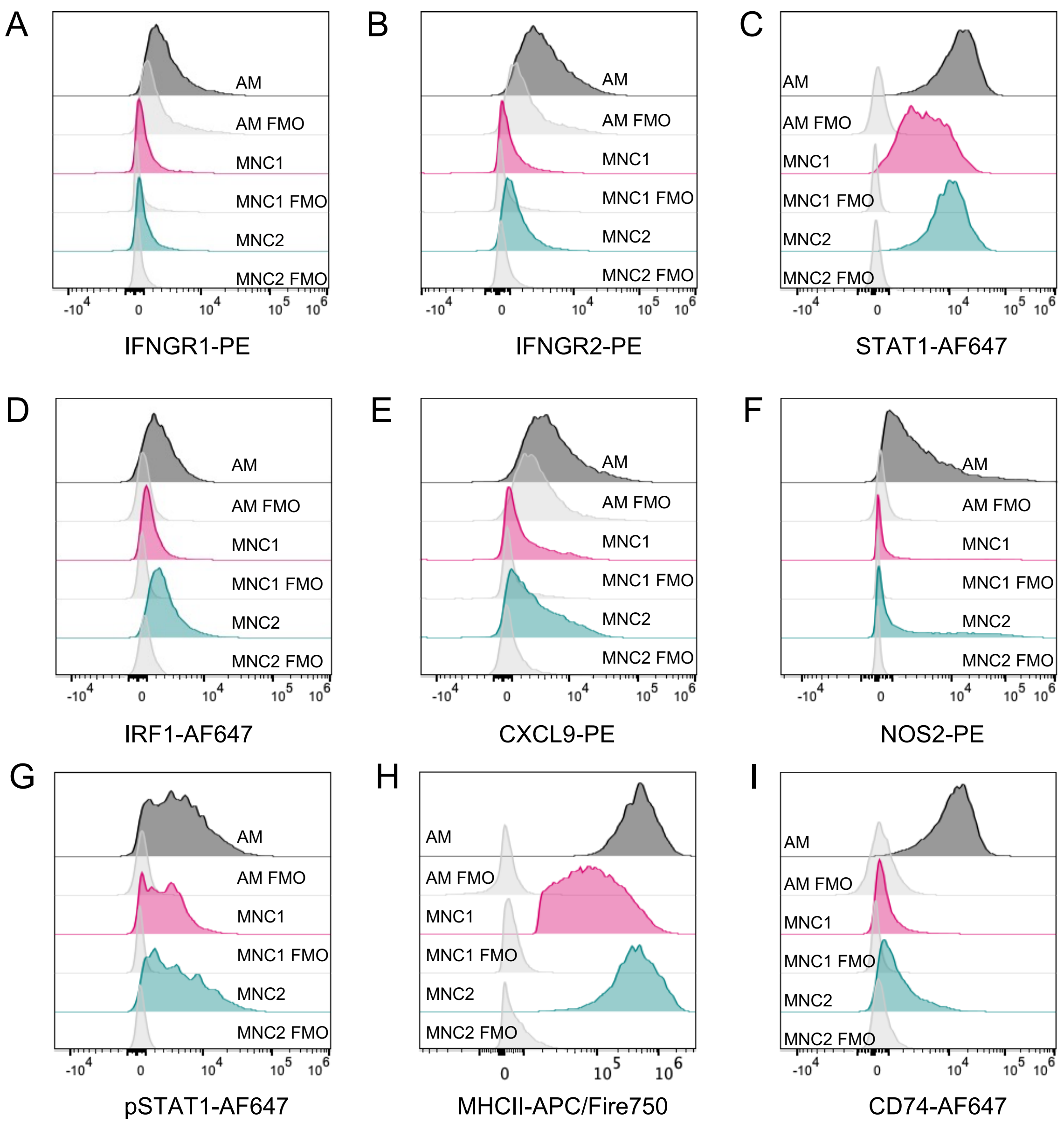


**Figure S2. Representative histograms of proteins analyzed by flow cytometry for lung MNP subsets.** FMO: fluorescence minus one.

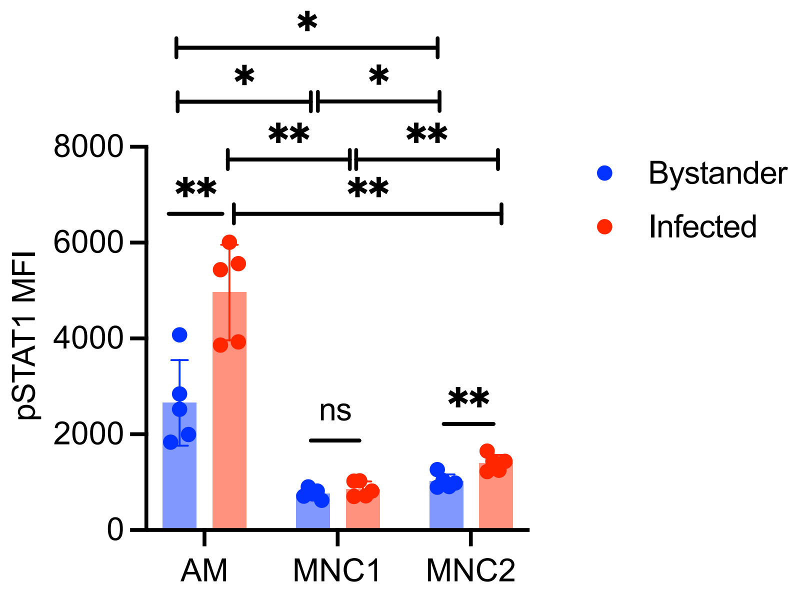


**Figure S3. Impaired IFN-γ responsiveness of MNC1 cells compared to other subsets in bystander or infected cells.**
C57BL/6 mice were infected with Mtb H37Rv-ZsGreen for 28 days. Lung single-cell suspensions were harvested and treated with 20 ng/mL IFN-γ for 15 min and subjected to pSTAT1 flow cytometry analysis. Results for bystander subsets ($ZsGreen^-$) and infected subsets ($ZsGreen^+$) are presented as mean ± SD. *p<0.05, **p<0.01 by Brown-Forsythe and Welch ANOVA tests with Dunnett's T3 multiple comparisons test (between subsets) or unpaired t-test (bystander versus infected subset); ns, not significant.

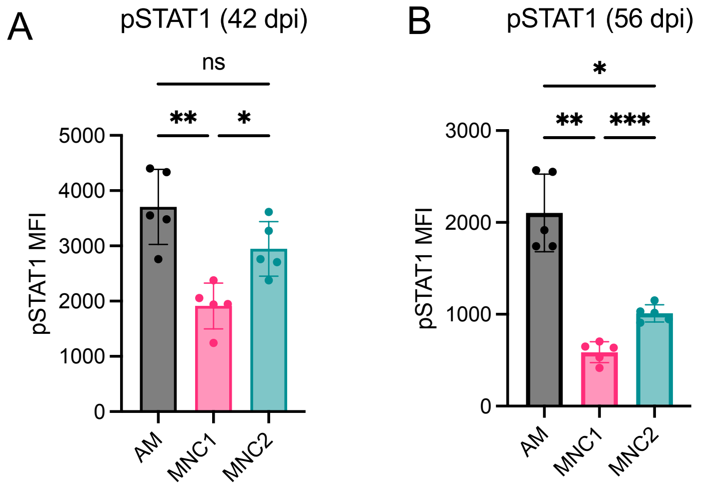


**Figure S4. MNC1 cells are less responsive to IFN-γ at multiple time points of chronic Mtb infection.**
C57BL/6 mice were infected with Mtb H37Rv-ZsGreen for 42 days (A) or 56 days (B). Lung single-cell suspensions were incubated with 20 ng/mL IFN-γ for 15 min and subjected to pSTAT1 flow cytometry analysis. Results are presented as mean ± SD.

**p<0.01, ***p<0.001, ****p<0.0001 by Brown-Forsythe and Welch ANOVA tests with Dunnett's T3 multiple comparisons test; ns, not significant.

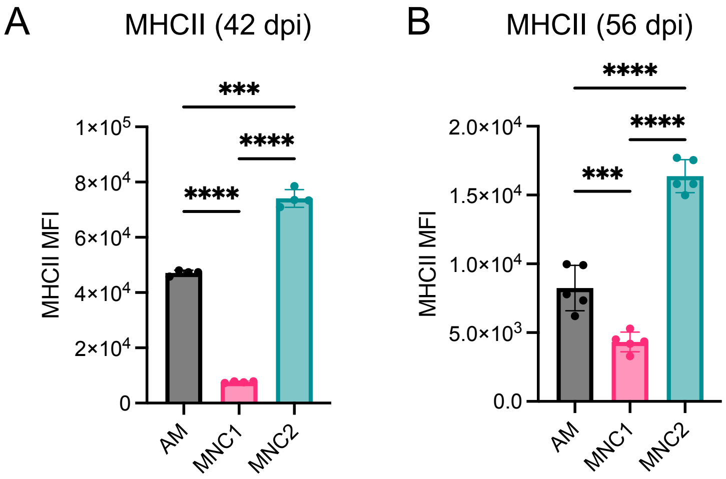


**Figure S5. MHC-II expression level in MNP subsets at multiple time points of chronic Mtb infection.** C57BL/6 mice were infected with Mtb H37Rv-ZsGreen for 42 days or 56 days. Lung cells were harvested for flow cytometry analysis. Results are shown as mean ± SD. ***p<0.001, ****p<0.0001 by Brown-Forsythe and Welch ANOVA tests with Dunnett's T3 multiple comparisons test.

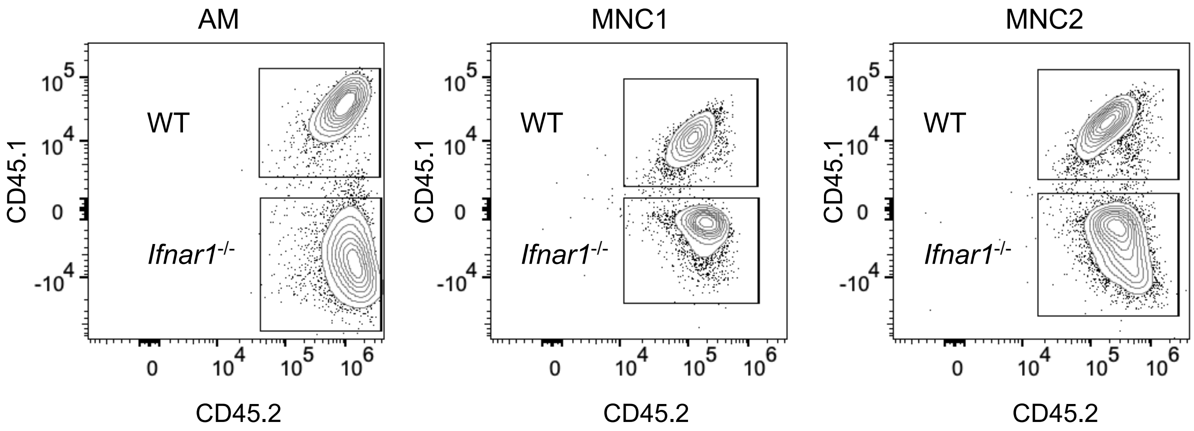


**Figure S6. Gating strategy for WT and *Ifnar1$^{-/-}$* subsets from mixed bone marrow chimeric mice.** Gating of AM, MNC1, and MNC2 is shown in Figure S1.

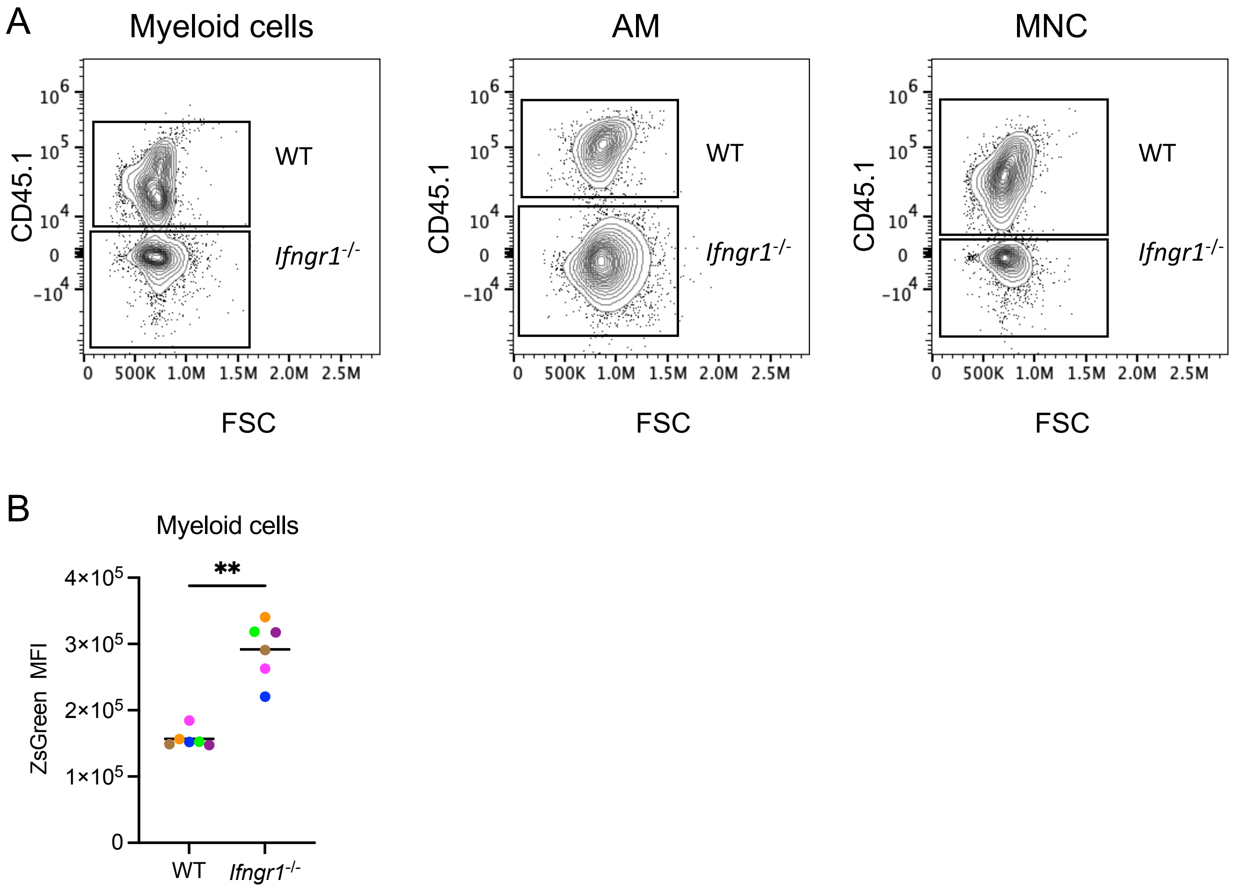


**Figure S7. Gating strategy and Mtb burden for WT and *Ifngr1$^{-/-}$* subsets.**
(A) Gating strategy for WT and *Ifngr1$^{-/-}$* subsets. Gating of myeloid cells, AM, and MNC is shown in Figure S1.
(B) Mtb burden for WT and *Ifngr1$^{-/-}$* myeloid cells.

### Table S1. IFN-γ responses differ in MNC1 and other MNP subsets

| Category | # of DEG in category | # of genes in category | term | p-value | Adjusted p-value | MNC1 vs |
|---|---|---|---|---|---|---|
| GO:0034341 | 64 | 135 | response to interferon-gamma | 1.16E-05 | 4.36E-02 | AM |
| GO:0034341 | 70 | 135 | response to interferon-gamma | 2.29E-13 | 1.09E-09 | MMC2 |
| GO:0071346 | 56 | 108 | cellular response to interferon-gamma | 9.33E-11 | 4.32E-07 | MMC2 |

### Table S2. MHC-II Antigen presentation pathways differ in MNC1 and other MNP subsets

| Category | # of DEG in category | # of genes in category | term | p-value | Adjusted p-value | MNC1 vs |
|---|---|---|---|---|---|---|
| GO:0002495 | 17 | 21 | antigen processing and presentation of peptide antigen via MHC class II | 5.63E-07 | 2.24E-03 | AM |
| GO:0002495 | 16 | 21 | antigen processing and presentation of peptide antigen via MHC class II | 1.00E-07 | 4.35E-04 | MMC2 |
| GO:0002504 | 19 | 23 | antigen processing and presentation of peptide or polysaccharide antigen via MHC class II | 9.15E-08 | 3.74E-04 | AM |
| GO:0002504 | 17 | 23 | antigen processing and presentation of peptide or polysaccharide antigen via MHC class II | 1.21E-07 | 5.25E-04 | MMC2 |
| GO:0019886 | 15 | 19 | antigen processing and presentation of exogenous peptide antigen via MHC class II | 4.85E-06 | 1.86E-02 | AM |
| GO:0019886 | 14 | 19 | antigen processing and presentation of exogenous peptide antigen via MHC class II | 1.28E-06 | 5.34E-03 | MMC2 |

### Table S3. Antibodies for flow cytometry

| Antibodies | Source | Identifier |
|---|---|---|
| Brilliant Violet 421 anti-mouse Ly-6G Antibody | BioLegend | Cat#127627; RRID: AB_10897944 |
| Brilliant Violet 605 anti-mouse CD11c Antibody | BioLegend | Cat#117334; RRID: AB_2562415 |
| Brilliant Violet 711 anti-mouse/human CD11b Antibody | BioLegend | Cat#101242; RRID: AB_2563310 |
| PE/Cyanine5 anti-mouse CD90.2 (Thy1.2) Antibody | BioLegend | Cat#105314; RRID: AB_313185 |
| PE/Cyanine5 anti-mouse CD19 Antibody | BioLegend | Cat#115510; RRID: AB_313645 |
| PE/Cyanine5 anti-mouse NK-1.1 Antibody | BioLegend | Cat#108716; RRID: AB_493590 |
| PE/Cyanine7 anti-mouse Ly-6C Antibody | BioLegend | Cat#128018; RRID: AB_1732082 |
| PE anti-mouse I-A/I-E Antibody | BioLegend | Cat#107608; RRID: AB_313323 |
| Alexa Fluor® 647 anti-mouse I-A/I-E Antibody | BioLegend | Cat#107618; RRID: AB_493525 |
| APC/Fire™ 750 anti-mouse I-A/I-E Antibody | BioLegend | Cat#107652; RRID: AB_2616729 |
| PE Rat Anti-Mouse Siglec-F | BD Biosciences | Cat#552126; RRID: AB_394341 |
| Alexa Fluor® 647 Rat Anti-Mouse Siglec-F | BD Biosciences | Cat#562680; RRID: AB_2687570 |
| CXCL9 (MIG) Monoclonal Antibody (MIG-2F5.5), PE, | eBioscience | Cat# 12-3009-80; RRID: AB_891582 |
| IRF-1 (D5E4) Rabbit mAb, Alexa Fluor 647 | Cell Signaling Technology | Cat# 14105S; RRID: AB_2798393 |
| Phospho-Stat1 (Tyr701) (58D6) Rabbit mAb, Alexa Fluor 647 | Cell Signaling Technology | Cat# 8009S; RRID: AB_10860764 |
| Alexa Fluor 647 Mouse anti-Total Stat1 (N-Terminus) | BD Biosciences | Cat#558560; RRID: AB_647143 |
| Purified Rat Anti-Mouse CD16/CD32 (Mouse BD Fc Block™) | BD Biosciences | Cat#553142; RRID: AB_394656 |
| PE anti-mouse IFNGR2 Antibody | Miltenyi Biotec | Cat# 130-105-671; RRID:AB_2652252 |
| PE anti-mouse IFNGR2 Antibody | BioLegend | Cat# 113604; RRID: AB_313561 |
| CD119 (IFN gamma Receptor 1) Monoclonal Antibody (2E2), PE | Invitrogen | Cat# 12-1191-82; RRID:AB_1210730 |
| iNOS Monoclonal Antibody (CXNFT), PE | eBioscience | Cat# 12-5920-82; RRID: AB_2572642 |
